# A somatic multiple myeloma mutation unravels a mechanism of oligomerization-mediated product inhibition in GGPPS

**DOI:** 10.1101/2024.12.25.628878

**Authors:** Ruba Yehia, Jasmína Mária Portašiková, Rut Mor Yosef, Benny Da’adoosh, Alan Kádek, Petr Man, Moshe Giladi, Yoni Haitin

**Affiliations:** Department of Physiology and Pharmacology, Faculty of Medical and Health Sciences, Tel-Aviv University, Tel-Aviv, 6997801, Israel; Institute of Microbiology of the Czech Academy of Sciences, Division BioCeV, Prumyslova 595, 252 50 Vestec, Czech Republic; Department of Biochemistry, Faculty of Science, Charles University, Hlavova 2030/8, 128 43, Prague 2, Czech Republic; BLAVATNIK CENTER for Drug Discovery, Tel Aviv University, Tel Aviv, Israel; Tel Aviv Sourasky Medical Center, Tel Aviv, 6423906, Israel; Sagol School of Neuroscience, Tel Aviv University, Tel Aviv, 6997801, Israel

## Abstract

Protein prenylation regulates the cellular localization of small GTPases and is pivotal for multiple myeloma (MM) pathology. Geranylgeranyl diphosphate synthase (GGPPS), synthesizing a prenylation moiety, exhibits dimeric or hexameric stoichiometry in different species. However, the functional significance of this divergence remains elusive. Focusing on the hexameric human paralog, formed by trimer-of-dimers, we uncover that GGPPS^R235C^, expressed in an MM cell line, localizes to the active site lid region at the inter-dimeric interface. Using crystallography and mass spectrometry (MS), we show that GGPPS^R235C^ retains its hexameric stoichiometry but exhibits destabilized inter-dimer interactions. Unexpectedly, this results in increased apparent substrate affinity and product release kinetics. These functional effects are further enhanced in a dimeric mutant, GGPPS^Y246D^. Combining MS and fluorescence spectroscopy, we exposed that reduced lid dynamics and increased active site occupancy by the product are intertwined. Together, our results expose product inhibition as a regulatory mechanism in GGPPS, driven by hexamerization.

## Introduction

Protein prenylation is a common post-translational modification (PTM) shared by 2% of the proteome (*1*), playing important roles in cellular function and survival, and occurs in all eukaryotic organisms. Prenylation contributes to protein localization by enabling their membrane association, allowing downstream cellular processes such as differentiation, proliferation, and trafficking to occur (*2*, *3*). Moreover, prenylation contributes to protein-protein interactions and facilitates signal transduction, among other roles (*4*).

During prenylation, the farnesyl or geranylgeranyl moiety of farnesyl diphosphate (C_15_; FPP) or geranylgeranyl diphosphate (C_20_; GGPP), respectively, is attached to a CaaX motif (where C is cysteine, a is an aliphatic amino acid and X is any amino acid, usually serine, glutamine or methionine) on the C-terminus of the target proteins, making it more hydrophobic and enabling its attachment to the membrane (*5*). Prenylated proteins include small GTPases, such as Ras, Rho, and Rab (*2*, *6*). These proteins are involved in a plethora of cellular processes, such as apoptosis, cytoskeleton formation, and cell cycle progression, and therefore are tightly regulated (*7*). Indeed, alterations in small GTPases’ function, protein-protein interactions, and the extent of prenylation result in malignant changes responsible for cancerous transformation and progression (*8*).

Geranylgeranyl diphosphate synthase (GGPPS) is a *trans*-prenyltransferase (PT) synthesizing GGPP by the condensation of FPP with isopentenyl diphosphate (C_5_; IPP) (*5*). GGPPS exhibits the canonical topology of *trans*-PTs, composed of 13 α-helices, forming dimers through a 4-layer helix bundle. 10 of the 13 α-helices surround a large central cavity harboring two conserved DDxxD motifs called aspartate-rich motifs (ARM) (*9*, *10*). Intriguingly, while sharing a similar topology, GGPPS orthologs display varying stoichiometries. Indeed, while most *trans*-PTs feature frequently-observed homodimeric assemblies (*11*), the pool of GGPPS oligomerization states is vast (*12*). Lower-level species predominantly exhibit the dimeric form of GGPPS, with some bacterial and plant species also displaying tetrameric and hexameric forms. In higher-level species, such as humans, GGPPS is a hexamer composed of a trimer of dimers (*5*, *11*). The N-terminal part of each monomer participates in both hexamer and dimer formation, with the inter-dimeric interface maintained by hydrophobic and H-bond interactions (*5*, *8*). Notably, despite the species-related stoichiometry deviation, the role of the different oligomeric compositions remains elusive.

With the pivotal role of geranylgeranylation in regulating the localization and activity of small GTPases, GGPPS has long been considered a viable target for novel anti-cancerous treatments (*12*). A prominent example is Multiple myeloma (MM), a malignancy of antibody-producing clonal plasma cells (*13*, *14*). In MM, Rab proteins are known as crucial for antibody production (*15–17*), and GGPPS inhibition was shown to induce the apoptosis of MM cells by disrupting Rab geranylgeranylation (*18*).

One of the commonly used cell lines in MM research (RPMI-8226) (*19*), previously shown to be sensitive to GGPPS inhibition (*20–22*), was reported to contain a somatic missense mutation in GGPPS (*23*) (Fig. 1A). Intriguingly, this mutation, GGPPS^R235C^, is localized to the inter-dimeric interface. Due to the central role of geranylgeranylation in MM pathophysiology and the sensitivity of RPMI-8226 cells to GGPPS inhibition, we explored the possible impact of the R235C mutation on the stoichiometry and activity of GGPPS here. Using functional, structural, and mass-spectrometry analyses, we expose a regulatory role for GGPPS hexamerization, mediated by the stabilization of inhibitory product binding within the active site. These findings shed light on the role of oligomerization in prenyltransferase evolution and provide structural insights to facilitate the design of novel GGPPS inhibitors.

**Fig. 1.**
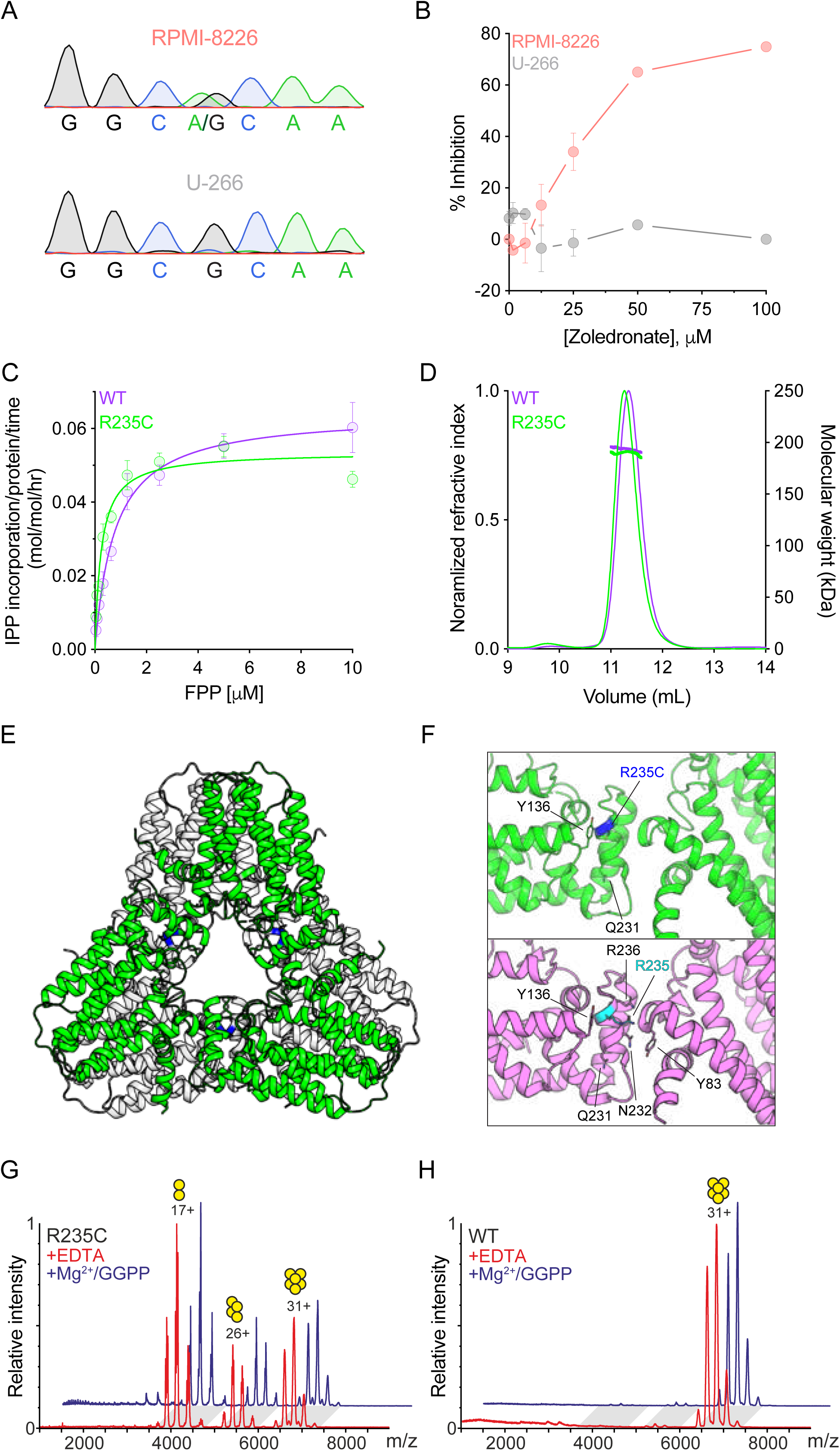
Identification and characterization of GGPPS^R235C^ from an MM model cell line. **(A)** Sanger sequencing of RPMI-8226 cells shows heterozygosity for the R235C mutation (left). This mutation is not present in U-266 cells (right). **(B)** Zoledronate-induced proliferation inhibition of the indicated cell lines, as measured using the XTT assay. **(C)** FPP-dependent activity of GGPPS^WT^ and GGPPS^R235C^. Data are shown as mean ± SEM, n = 3. **(D)** Representative SEC-MALS analysis. The black and green thin curves indicate the normalized refractive index of GGPPS^WT^ and GGPPS^R235C^, respectively. MALS-derived molecular weights are presented as bold horizontal lines. **(E)** Crystal structure of GGPPS^R235C^. The asymmetric unit, containing the biological hexameric arrangement, is shown in a cartoon representation. For spatial clarity, within each dimer, the subunits are differentially colored in grey and green. **(F)** Blowout on the inter-dimer interface in GGPPS^R235C^ (top) and GGPPS^WT^ (bottom). **(G, H)** nESI-MS spectrum obtained using low activation conditions (10 V trap collision voltage) for GGPPS^R235C^ (G) and GGPPS^WT^ (H). While GGPPS^R235C^ exhibits m/z distributions corresponding to several oligomeric states, GGPPS^WT^ is uniformly hexameric (yellow circles).

## Results

### GGPPS^R235C^ exhibits increased catalytic activity

Metabolic pathway addiction, supporting dysregulated cellular proliferation, is a cancer hallmark (*24*, *25*). With the suggested central role of GGPPS in MM biology (*12*), we explored the COSMIC database (*23*) for functionally significant cancer-associated mutations in GGPPS. Surprisingly, a GGPPS mutant is reported in RPMI-8226 (*26*), an MM cell line commonly used for drug discovery endeavors (*19*). Sanger sequencing revealed that these cells are heterozygous for the non-conservative GGPPS^R235C^ mutation, while this mutation was absent from U-266, another common MM cell line (Fig. 1A). Interestingly, these cell lines show markedly different responses to incubation with the known GGPPS inhibitor, zoledronate (*27*) (Fig. 1B). While U-266 cells are resistant to zoledronate, RPMI-8226 show clear dose-dependent inhibition of proliferation, supporting the apparent functional significance of GGPPS^R235C^ in these cells (*28*). Next, to determine the functional effect of GGPPS^R235C^, we proceeded with enzyme kinetics characterization of the purified enzyme (Fig. 1C). Surprisingly, while the mutant exhibited a similar *k*_cat_ to the WT (0.05 ± 0.003 s^-1^ vs. 0.06 ± 0.003 s^-1^, respectively), the K_m_ for FPP was four-fold reduced (0.2 ± 0.05 μM vs. 0.8 ± 0.15 μM, respectively).

As R235C is positioned at helix α10, which forms part of the inter-dimer interface, we next sought to determine its possible effect on GGPPS oligomeric state (Fig. 1D). SEC-MALS analysis revealed that GGPPS^R235C^ maintains its hexameric stoichiometry in solution, indicating that the reduced K_m_ does not result from the disintegration of the complex. Moreover, crystallographic analysis of GGPPS^R235C^ did not reveal significant global structural alterations (Fig. 1E) nor local changes at the interdimeric interface (Fig. 1F). Similar to the structure of GGPPS^WT^ (*5*), despite carrying our purification and crystallization without GGPP addition, we identified a GGPP molecule within the active site of each monomer (Supplementary Fig. 1). These molecules, bound at the suggested inhibitory product position (*5*), originate from the expression system itself and suggest that highly tight interactions of GGPP result in product inhibition. Notably, analysis of this structure using the PDBe PISA server (*29*) revealed a reduction in the predicted stability of the complex, with ι1G^diss^ of 16.9 kcal/mol and 20.1 kcal/mol for the GGPP^R235C^ and GGPP^WT^ proteins, respectively.

To further explore the impact of GGPPS^R235C^ on the integrity of the complex, we next performed native electrospray ionization mass-spectrometry (nESI-MS) analysis. Consistent with the predicted reduction in complex stability, as reflected by ι1G^diss^, we observed the dissociation of GGPPS^R235C^ (Fig. 1G), but not the WT proteins (Fig. 1H), into tetramers and dimers already at very low activation energy (10 V trap collision voltage). This behavior was not affected by the addition of EDTA to chelate substrate-coordinating Mg^2+^ ions or adding saturating concentrations of Mg^2+^ and GGPP. Thus, GGPPS^R235C^ weakens the inter-dimeric interface, thereby destabilizing the hexameric organization of the complex.

### The inter-dimeric interface regulates enzymatic function

GGPPS^R235C^ exhibits reduced K_m_ (Fig. 1C) along with a reduction in the hexameric stability (Fig. 1G,H). In order to establish the possible structure-function correlate of these observations, we took advantage of the GGPPS^Y246D^, a mutant previously reported to exhibit dimeric stoichiometry (*19*). In agreement with previous studies, SEC-MALS analysis of GGPPS^Y246D^ demonstrates an overwhelming dimeric population in solution (Fig. 2A). Moreover, nESI-MS corroborates the presence of stable dimers in a manner independent of active site occupancy by Mg^2+^ ions or GGPP (Fig. 2B). Strikingly, the GGPPS^Y246D^ mutation resulted in significantly increased *k*_cat_ to the WT (0.15 ± 0.006 s^-1^ vs. 0.06 ± 0.003 s^-1^, respectively), with a four-fold reduction in the K_m_ for FPP (0.2 ± 0.04 μM vs. 0.8 ± 0.15 μM, respectively), similarly as in the R235C mutant (Fig. 2C). To determine the possible effect of interrupting the inter-dimeric interface on active site architecture, we determined the crystal structure of GGPPS^Y246D^. In contrast with GGPPS^WT^ and GGPPS^R235C^, GGPPS^Y246D^ crystalized in the apo state (Fig. 2D). Importantly, comparison of the two GGPPS subunits revealed two different conformations of helices α10 and α11 (Fig. 2E), forming a ‘lid’ over the active site and contributing to the inter-dimer interface. Specifically, while the ‘lid’ conformation in one subunit is structurally preserved, displaying a reminiscent fold to that observed in GGPPS^WT^, the other subunit shows a markedly different conformation. This perturbation results, in turn, in a significant widening of the passageway between the active site and surrounding bulk due to the retraction of helix α9a at the mouth of the active site, increasing its distance from α3-α4 linker (S70-E206, Cβ-Cβ) from 5.6 Å to 7.2 Å and 12 Å for chains B and A, respectively. Interestingly, GGPPS^R235C^ displays a WT-like structural organization, in line with its preserved oligomeric assembly and *k*_cat_ (Fig. 2F).

**Fig. 2.**
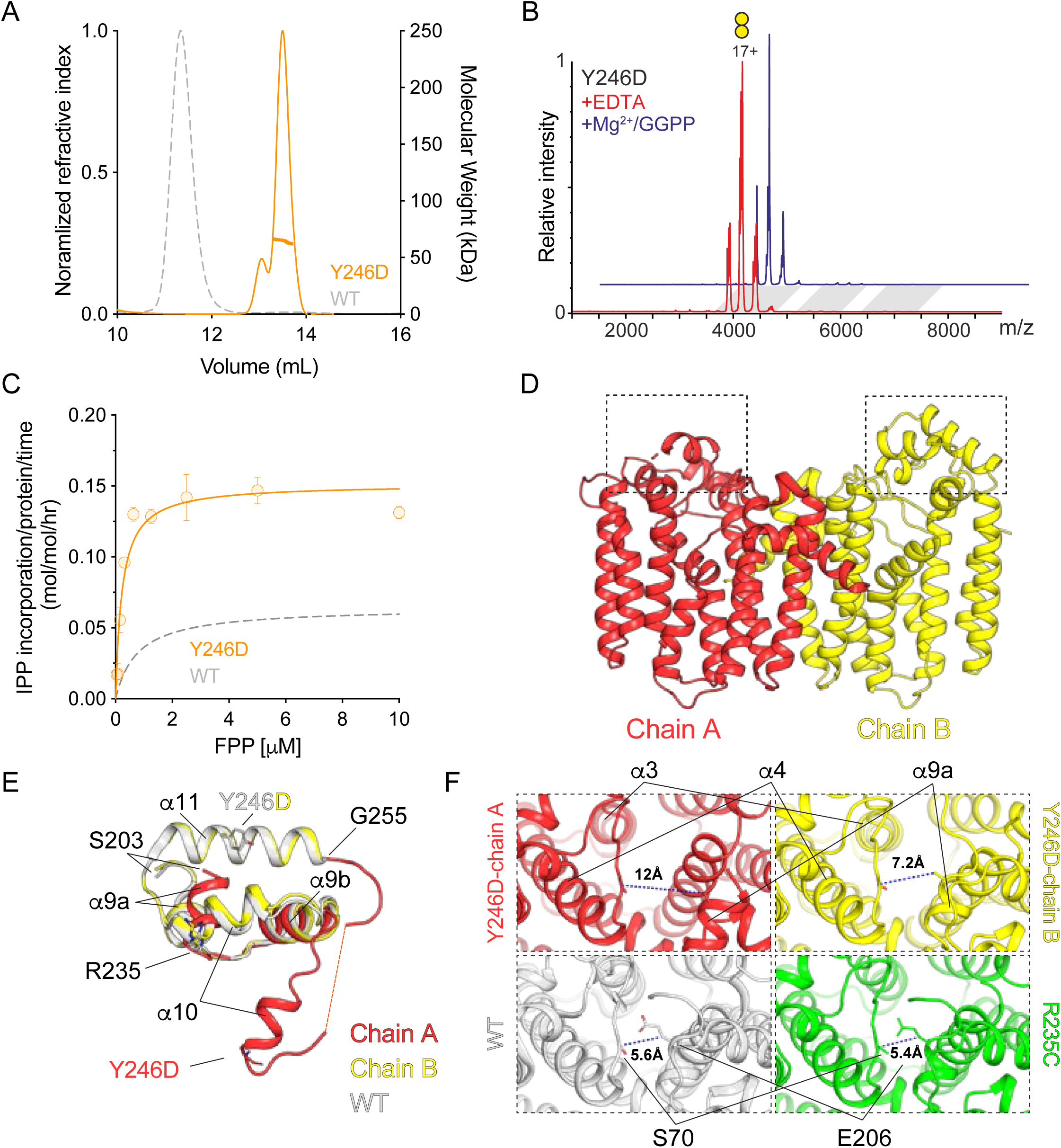
Structural and functional analyses of the dimeric GGPPS^Y246D^. **(A)** Representative SEC-MALS analysis. The orange curve indicates the normalized refractive index of GGPPS^Y246D^. The MALS-derived molecular weight is presented as a bold horizontal line. For reference, the SEC profile of GGPPS^WT^ is shown in grey. **(B)** nESI-MS spectrum obtained using low activation conditions (10 V trap collision voltage) for GGPPS^Y246D^, displaying a uniform dimeric stoichiometry (yellow circles). **(C)** FPP-dependent activity of GGPPS^Y246D^. and Data are shown as mean ± SEM, n = 3. For reference, the enzyme kinetics curve of GGPPS^WT^ is shown in grey. **(D)** Crystal structure of GGPPS^Y246D^. The asymmetric unit, containing a single dimer, is shown in a cartoon representation. For spatial clarity, within each dimer, the subunits are differentially colored in red and yellow. **(E)** Superposition of the lid region, framed in a dashed rectangle in (D) between each subunit and GGPPS^WT^. **(F)** A comparison of the lid region arrangement in the different GGPPS structures reveals a larger opening in GGPPS^Y246D^.

### The Inter-dimeric interface contributes to product inhibition

Previous crystallographic studies of hGGPPS, unless incubated with the known bisphosphonate inhibitors (*27*), revealed a GGPP molecule bound at an active site position considered inhibitory (Supplementary Figs. 1 and 2) (*5*). This is in contrast to previously reported dimeric GGPPS homologs from lower-level species (*30*) or due to the introduction of disease-causing mutation to the active site of hGGPPS (*31*), where apo structures have been reported. In line with the structural data, nESI analysis of hGGPPS^WT^ revealed that the active site is substantially occupied by GGPP, even in the absence of magnesium ions, which participate in GGPP coordination (Fig. 3A, Supplementary Fig. 1). Specifically, hGGPPS^WT^ hexamers exhibit Gaussian-like mass distribution, with three and four GGPP-bound subunits most abundant. Notably, incubation of the enzyme with saturating magnesium and GGPP concentrations results in a shifted mass distribution towards fully GGPP-occupied complexes (Fig. 3A). These results may reflect a dynamic equilibrium with a slow release of GGPP from the active site of hGGPPS. Intriguingly, while hGGPPS^R235C^ exhibits a range of occupancy states, reminiscent of the WT complex, the addition of magnesium and GGPP produces a milder shift in the occupancy distribution (Fig. 3B). This observation may provide the explanation for the lower K_m_ observed in the enzyme kinetics assay (Fig. 1C), stemming from the reduction in competitive product inhibition. Finally, hGGPPS^Y246D^ also displays occupancy distribution, albeit with a significant bias towards the apo state (Fig. 3C), as supported by our crystallographic analysis (Fig. 2D).

**Fig. 3.**
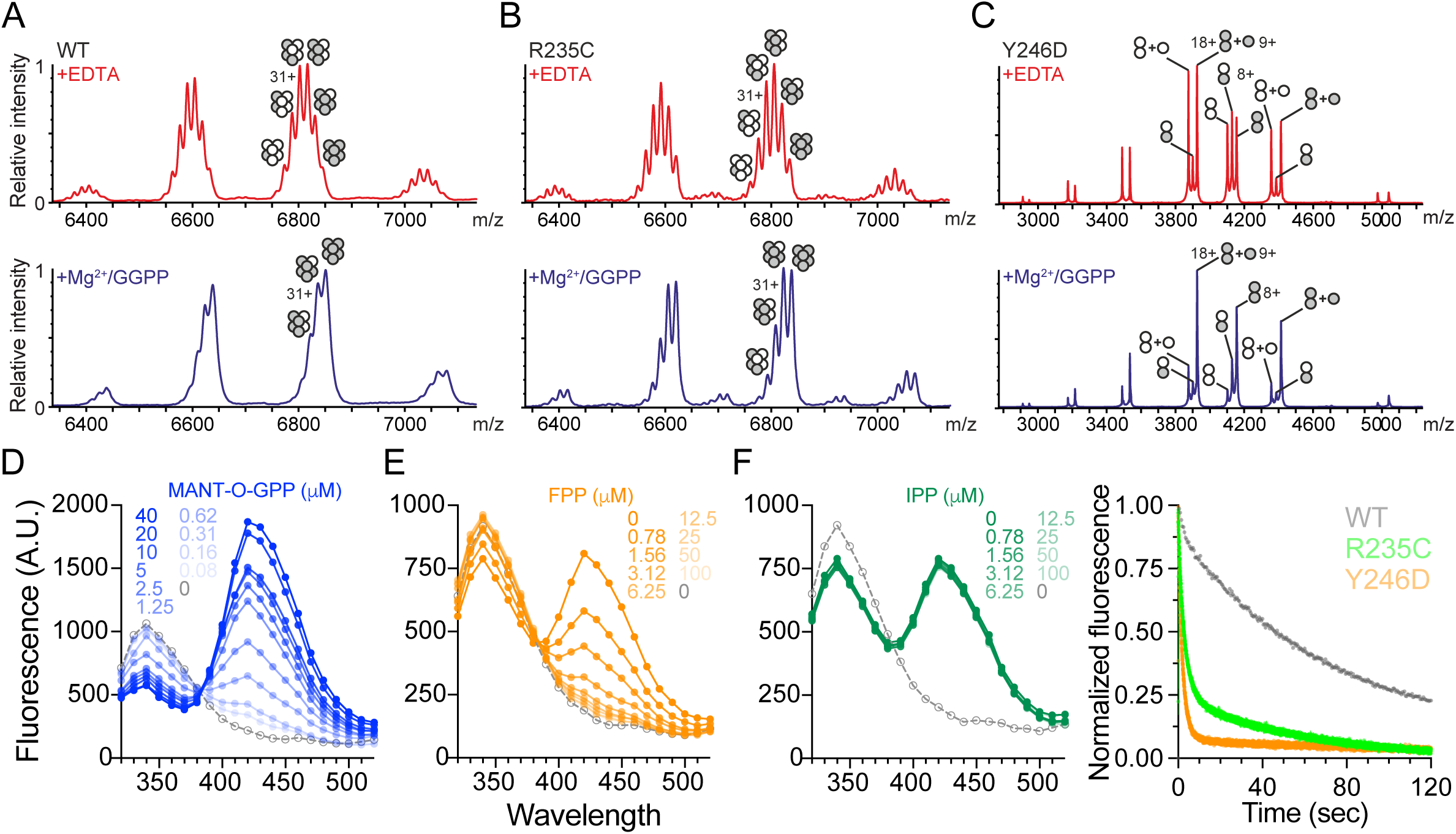
Inter-dimer interface mutations affect active site occupancy. **(A-C)** nESI-MS spectrum obtained using high activation conditions (80 V trap collision voltage) for GGPPS^WT^ (A), GGPPS^R235C^ (B), and GGPPS^Y246D^ (C). The occupancy states of the active sites are indicated above the peaks as grey circles. **(D-F)** Spectral FRET analysis of MANT-O-GPP interaction with GGPPS. Average fluorescence emission spectra following excitation at *F*_280_ (n = 3). Increasing concentrations of MANT-O-GPP resulted in a concomitant decrease of *F*_340_ and an increase of *F*_420_ in GGPPS^WT^ (D). Incubation with increasing concentrations of FPP (E) but not IPP (F) reverted this spectral shift, indicating that MANT-O-GPP occupies the FPP/GGPP site. **(G)** Averaged stopped-flow kinetics of MANT-O-GPP displacement from GGPPS^WT^, GGPPS^R235C^, and GGPPS^Y246D^, measured using FRET (n = 5).

To determine if the observed perturbation of the dynamic equilibrium reflects altered product release kinetics, we resorted to a combined FRET-based stopped-flow approach (*32*) (Fig. 3D-G). To this end, we used the fluorescent GGPP analog (2E,6E)-8O-(N-methyl-2-aminobenzoyl)-3,7-dimethyl-2,6-octandien-1pyrophosphate (MANT-O-GPP) (*33*, *34*). As the excitation profile of MANT-O-GPP closely resembles the emission profile of tryptophan (*32*), it can serve as FRET acceptor for native GGPPS tryptophan donors, thereby reporting on active site occupancy. Indeed, a dose-response increase in the FRET signal is observed with increasing MANT-O-GPP concentrations (Fig. 3D). Notably, steady-state FRET-based competition experiments indicated that MANT-O-GPP could be displaced by FPP (Fig. 3E) but not IPP (Fig. 3F). Taken together with previous structural analyses, MANT-O-GPP exhibits a binding mode reminiscent to that of the inhibitory GGPP (Supplementary Fig. 2). To monitor the displacement of MANT-O-GPP from the inhibitory site, GGPPS was incubated with excess MANT-O-GPP, followed by rapid mixing with a 20-fold molar excess of FPP, to prevent MANT-O-GPP rebinding, resulting in FRET signal decrease (Fig. 3G). In agreement with the crystallographic and nESI results, MANT-O-GPP dissociates from hGGPPS^WT^ with markedly slow biphasic kinetics (*k*_fast_ = 0.02 ± 0.0004 s^-1^ and *k*_slow_ = 0.003 ± 0.00005 s^-1^). Remarkably, hGGPPS^R235C^ displayed significantly accelerated dissociation kinetics (*k*_fast_ = 0.4 ± 0.001 s^-1^ and *k*_slow_ = 0.03 ± 0.0008 s^-1^). The effect was even greater for hGGPPS^Y246D^, which exhibited monophasic dissociation kinetics (*k*_fast_ = 0.5 ± 0.003 s^-1^). These results suggest that destabilization of the inter-dimer interface enables faster dissociation of the inhibitory product, resulting in increased catalytic activity.

### The Inter-dimeric interface restricts the active site ‘lid’ dynamics

In order to explore the possible correlation between substrate dissociation and the stability of the inter-dimeric interface, we used hydrogen-deuterium exchange mass spectrometry (HDX-MS) analysis of GGPPS^WT^ and its mutants (Fig. 4). Following time-resolved incubation in a reaction buffer made of D_2_O and subsequent proteolytic fragmentation, the rate of backbone HDX of each peptide is determined, providing spatially resolved information on structural dynamics (*35*). As the HDX rate depends on both solvent accessibility and the local state of hydrogen bonding and dynamics, the protein core, secondary structure elements, and buried interfacial regions shall exhibit lower HDX.

**Fig. 4.**
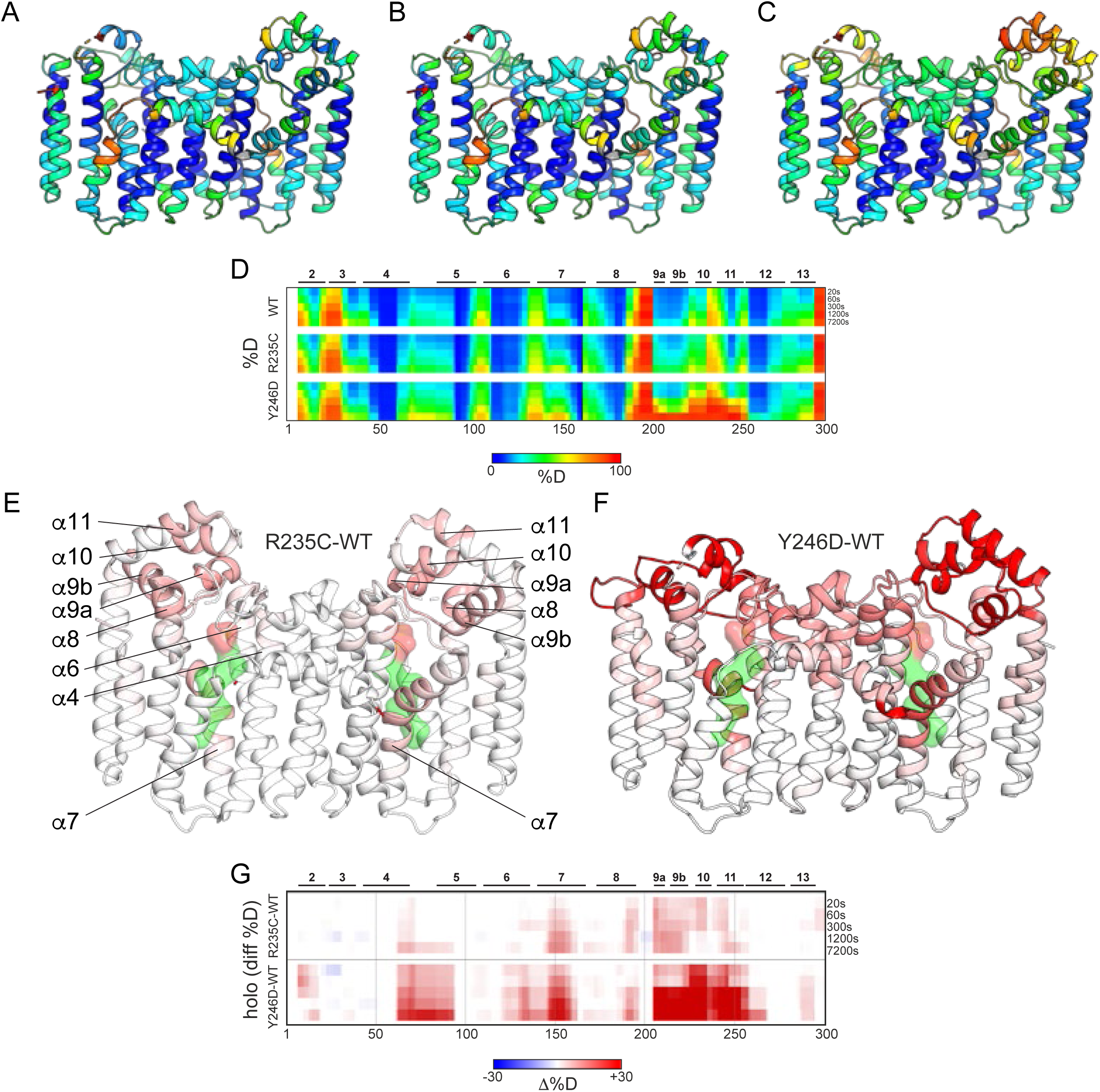
HDX-MS analyses support mutation-induced enhanced lid dynamics. **(A-C)** Representative dimers of GGPPS^WT^ (A), GGPPS^R235C^ (B), and GGPPS^Y246D^ (C). Heatmap coloring indicates the HDX level following 300 s incubation in D_2_O. The level of exchange is indicated in the provided color bar. **(D)** Deuteration levels at the indicated time points for the different GGPPS constructs. Secondary structure elements are indicated above the heat maps. **(E, F)** The HDX differences between GGPPS^R235C^ (E) or GGPPS^Y246D^ (F) and GGPPS^WT^ were mapped onto the structures of the corresponding mutants. Secondary structure elements are indicated. Surface representations of the GGPP molecules [placed by superposition with PDB 2Q80 (*5*)] are shown within the active site. **(G)** Heat maps summarizing differential HDX-MS analyses at the indicated time points of GGPPS^WT^ and its mutants. Secondary structure elements are indicated above the heat maps.

The HDX profiles were followed in the presence of a saturating concentration of GGPP and magnesium (Fig. 4A-D). As expected, GGPPS and its mutants exhibit very low HDX at the intra-dimer interface (helices α5 and α6; Fig. 4A-D). However, comparison of GGPPS^WT^ and the mutants demonstrated two main regions exhibiting varying degrees of increased deuteration rates (Fig. 4E-G) in both mutants, GGPPS^R235C^ (Fig. 4F) and GGPPS^Y246D^ (Fig. 4G). These are: (*1*) the ‘lid’ of the active site, and (*2*) the hydrophobic geranylgeranyl binding cavity. In both mutants, throughout the experimental time course, helices α8-α11 (the ‘lid’ region) exhibit increased deuterium uptake compared to the WT, reflecting enhanced exposure of the backbone to the bulk solution. Intriguingly, at the hydrophobic GGPP binding cavity, consisting of helices α4, α6, and α7, the HDX differences develop gradually over time (Fig. 4G). This is consistent with the ‘lid’ as the product release passageway, where product dissociation from the hydrophobic cavity is preceded by a conformational change at the ‘lid’ region. Indeed, in all regions, the differences in HDX are more prominent for GGPPS^Y246D^ than for GGPPS^R235C^ (Fig. 4F, G). Thus, the increased opening probability of the ‘lid’ region accelerates product release kinetics, thereby relieving product inhibition and increasing the catalytic activity.

## Discussion

Small GTPases are key players in cancer cell biology, underlying some of the hallmarks of cancer (*25*, *36*), including sustained proliferative signaling, resistance to cell death, induction of angiogenesis, and promotion of tissue invasion (*37*, *38*). Given the essential roles of small GTPases play in cellular survival, together with their functional dependence on membrane localization, prenylation inhibitors are considered a long-awaited therapeutic class (*38*, *39*).

Unfortunately, many of these compounds were found to be highly toxic, limiting their clinical value (*39*). In contrast, the specific inhibition of GGPPS was shown as a potential safe and effective therapeutic strategy for MM in cellular and animal models (*19*).

*In vitro* cancer models, though essential in anti-cancer drug development, present significant challenges due to their limited resemblance to actual clinical conditions. These discrepancies raise concerns about the reliability of findings derived from such models, as they may fail to capture the complexities of *in vivo* cancer biology (*40*). These limitations notwithstanding, we identified a missense mutation in GGPPS in RPMI-8226, a widely used cell line in MM drug discovery campaigns. Rummaging through the COSMIC database reveals dozens of missense mutations in GGPPS from different cancers. However, we found GGPPS^R235C^ particularly peculiar due to its strategic position at the inter-dimeric interface (Fig. 1E,F) and identification in an MM cell line (Fig. 1A). With the apparent increased sensitivity of these cells to GGPPS inhibition (Fig. 1B), we hypothesized that this mutation may affect the quaternary organization of the enzyme and, as a result, its function and contribution to the malignant phenotype of these cells (*41*).

Like numerous other oligomers (*42*), the hexameric human GGPPS shares a basic homodimeric functional unit with homologs scattered throughout the phylogenetic tree (*5*). Homooligomerization provides enzymes with structural and functional advantages crucial for their optimal performance in the cell (*43*). It can enhance enzyme stability by making it more resistant to environmental stresses such as temperature and pH fluctuations, facilitate allosteric regulation and cooperativity, and improve catalytic efficiency by bringing multiple active sites into proximity. Here, using multiple experimental approaches, we surprisingly expose GGPPS hexamerization as an intrinsic inhibitory mechanism, diminishing enzymatic function.

While GGPPS^R235C^ maintains its hexameric stoichiometry in solution, it displayed reduced stability. Specifically, nESI-MS experiments revealed that contrary to GGPPS^wt^, even during gentle ESI transfer into the gas phase, GGPPS^R235C^ undergoes dissociation into tetrameric and dimeric sub-assemblies, consistent with perturbation of the inter-dimeric interface (Fig. 1G). Unexpectedly, this reduced stability was accompanied by reduced K_m_ (Fig. 1C), indicating higher substrate affinity. To establish the relationship between the decreased stability of the inter-dimeric interface and the enhanced substrate affinity, we exploited the GGPPS^Y246D^ mutant, exhibiting purely dimeric stoichiometry. In line with our hypothesis, this mutant showed a similar decrease in K_m_ as GGPPS^R235C^ and an increase in catalytic activity (Fig. 2C). Considering the dichotomic segregation of GGPPS stoichiometry of GGPPS to hexameric and dimeric complexes throughout the phylogenetic tree, these results highlight the potential functional autoregulation by the quaternary organization of the enzyme.

Interestingly, controlled activation during the native MS analyses has also uncovered different stability of the dimeric versus higher oligomeric species of GGPPS. When exposed to energetic collisions with an inert gas (Ar) inside a mass spectrometer, hexameric GGPPS (both GGPPS^WT^ and GGPPS^R235C^) dissociates through the ejection of an unfolded monomer subunit as indicated by its disproportionately high charge state distribution (Supplementary Fig. 3A,B). On the other hand, dimeric species (both GGPPS^Y246D^ and GGPPS^R235C^) partition equally into lower charged and thus folded monomers (Supplementary Fig. 3D,E), while the GGPPS^R235C^ tetramer shows a mixture of both dissociation modes (Supplementary Fig. 3C). This suggests that the formation of higher oligomeric states conveys increased stability to the GGPPS assembly, which aligns well with the subunit interaction analysis of the crystal hexameric GGPPS^WT^ structure (PDB 2Q80) (*5*) by the PDBePISA service (*29*). The three intra-dimeric interfaces (e.g. between chains A-B) have large solvent-excluded contact interfaces with very few hydrogen bonds (on average 1579 Å^2^ with only 5 bonds). On the other hand, the six inter-dimeric contacts (e.g. between chains A-D and A-E), while about three times smaller in area size, bring substantially more hydrogen bonds (each on average 8 bonds on just 479 Å^2^), thus stabilizing the assembly through a chain of interactions between dimers. Therefore, it is tempting to speculate this may form a part of the activity modulation mechanism between different species in nature.

Beyond the differences in oligomeric assembly, structural analysis of GGPPS^R235C^ and GGPPS^Y246D^ exposes key differences in the architecture and occupancy of the active site. Similar to the hexameric GGPPS^WT^, GGPPS^R235C^ crystallized in complex with one GGPP molecule bound in an inhibitory configuration per monomer (Fig. 1E, Supplementary Fig. 1). In sharp contrast, GGPPS^Y246D^ crystallized in the apo state (Fig. 2D). Comparing the two structures, a large movement of the lid region is observed in GGPPS^Y246D^. The structures reveal that this region plays a dual role. On the one hand, it forms an integral part of the inter-dimer interface, stabilizing the hexameric organization of GGPPS. On the other hand, it is involved in the coordination of the pyrophosphate headgroup of GGPP in the inhibitory product configuration (Supplementary Fig. 2).

Product inhibition mechanisms are widespread, ensuring that the production rate of pathway end products aligns with their physiological need. In this mechanism, the end product itself acts as a regulator by inhibiting the activity of the enzyme, catalyzing its production. This inhibition prevents the overproduction of the end product when it is not being utilized, conserving essential cellular resources by avoiding the unnecessary production of unused compounds. Indeed, this form of regulation is both efficient and widespread throughout the kingdoms of life (*44*). Direct examination of the intact complex mass revealed lower active site product occupancy (Fig. 3A-C) due to markedly faster dissociation rates for both mutants, with GGPPS^Y246D^ displaying the greatest effect (Fig. 3G). Moreover, HDX-MS revealed that the lid region exhibits moderately increased dynamics in the context of the GGPPS^R235C^, further enhanced in GGPPS^Y246D^ (Fig. 4).

We suggest the following model for GGPPS product inhibition by GGPP (Fig. 5A). In the native hexameric complex, the lid region is sterically confined by the inter-dimer interface, priming it for a stable interaction with GGPP in the inhibitory conformation. Indeed, in this conformation, the strictly conserved K202, positioned with the α8-α9a loop, can directly interact with the inhibitory product, thereby establishing the lid as a bona fide component of the inhibitory site. With reduced stability of the inter-dimer interface, the inhibitory product release probability increases, as directly measured using stopped-flow. This, in turn, results in the alleviation of product inhibition and increased activity. Collectively, these results reinforce the correlation between lid flexibility and enzymatic function, providing a structural basis for GGPPS product inhibition by higher-order oligomerization (Fig. 5A).

**Fig. 5.**
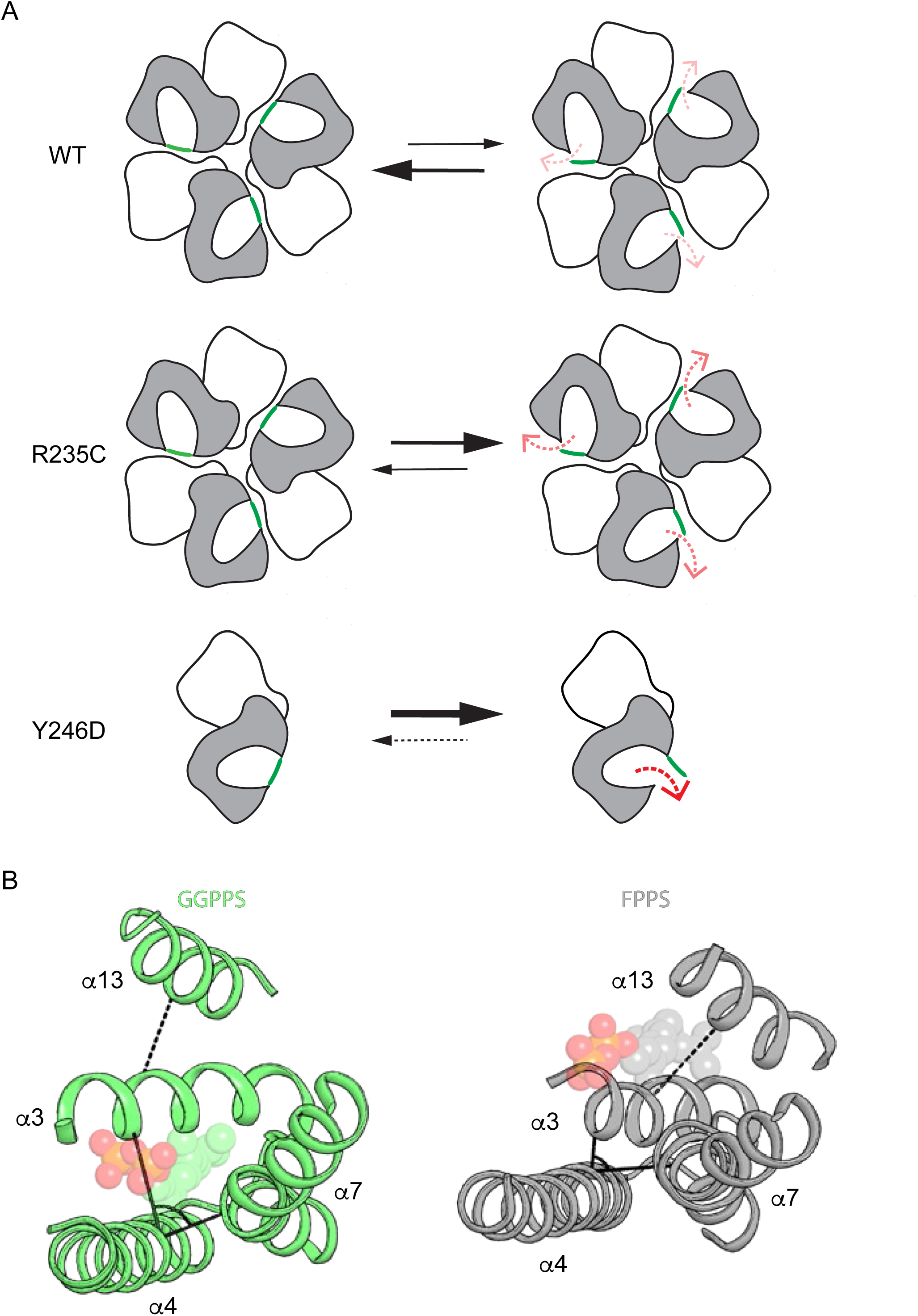
Proposed model for GGPPS product inhibition. **(A)** In GGPPS^WT^, the inter-dimer interface stabilizes the lid region in a closed state, resulting in a low product release probability. In GGPPS^R235C^, the hexameric stoichiometry is preserved, while the inter-dimer interface is destabilized, resulting in higher lid opening and product release probabilities. This effect is further exacerbated in the dimeric GGPPS^Y246D^ mutant. **(B)** Comparison of the product-inhibited state of GGPPS (PDB 2Q80) and FPPS (PDB 5JK0). In GGPPS, the cavity between α3, α4, and α7 accommodates the large GGPP product. Conversely, in FPPS, the FPP product resides between α3 and α13. GGPP and FPP are shown as semi-transparent spheres.

Product inhibition was previously reported for additional mevalonate pathway enzymes (*45*). This process was recently exposed in the human farnesyl pyrophosphate synthase (FPPS), an upstream branchpoint enzyme producing FPP (*46*). Intriguingly, despite their high structural similarity, FPPS and GGPPS achieve product inhibition via distinct structural mechanisms (Fig. 5B). Specifically, while GGPPS displays an inhibitory product binding site overlap with the active site cavity (Supplementary Fig. 2), FPPS shows an allosteric inhibitory site, confined by helices a3 and α13. The comparison between the crystal structures of both enzymes reveals that the active site of FPPS is unable to accommodate FPP in a conformation reminiscent of that of the inhibitory GGPP binding pose. This is related to the producrt-length determination mechanism, dictated by a smaller active site volume in FPPS (*47*). Conversely, the longer GGPP product cannot fit between helices α3 and α13 to satisfy the coordination of its polar headgroup, as seen in the case of FPP.

In conclusion, driven by the increased activity of a multiple myeloma-associated mutant, we exposed the functional role of hexamerization in GGPPS and the structural mechanism of product-mediated product inhibition. As GGPPS inhibitors present a long-sought therapeutic class, these results may facilitate the rational design of allosteric inhibitors targeting and stabilizing the inter-dimeric interface.

## Materials and Methods

### PCR and sequencing

RNA extraction from RPMI-8226 or U-266 cell was performed using MagMax mirVana RNA Isolation Kit (Thermo Fisher Scientific Waltham, MA, USA), followed by cDNA synthesis using Maxima H Minus RT (Thermo Fisher Scientific Waltham, MA, USA). The area flanking the R235C mutation on the *GGPS1* gene was amplified using PCR (Phusion high-fidelity PCR 2X master mix (New England Biolabs, MA, USA) with 10 μM of each primer). For the reaction, 2 ng/μl of the cDNA were amplified with two primers (forward: 5’- ATTGGTCAAGGCCTGAAAGCA-3′, reverse: 5’-TTCCCACCACGTGCATCAAT-3′), resulting in a 200 bp product, which embeds the R235C point mutation. Prior to sequencing with the forward primer mentioned above, the PCR product was purified using NucleoSpin Gel and PCR Clean-up (New England Biolabs, MA, USA).

### Cellular proliferation analysis

RPMI-8226 and U-266 cells were cultured in RPMI-1640 medium supplemented with 10% fetal bovine serum, 1% penicillin-streptomycin, and 2 mM L-glutamine at 37°C in a humidified 5% CO₂ incubator. Cellular proliferation was analyzed using the XTT assay (Sigma Aldrich, MO, USA) following the manufacturer’s protocol. Cells were seeded in 96-well plates at 1 × 10⁵ cells/well in 100 µL of medium and treated with 0-100 μM zoledronate. After 72 hours, 50 µL of XTT reagent mixture was added to each well and incubated for an additional 4 hours at 37°C. Absorbance was measured at 450 nm with a reference wavelength of 630 nm using a microplate reader. Proliferation was calculated as the percentage of absorbance in treated cells relative to untreated controls, with blank wells subtracted for background correction. Experiments were conducted in 4 replicates, with results expressed as mean ± SEM.

### Protein expression and purification

cDNA of full-length human *GGPS1* (transOMIC, Huntsville, AL, USA) was cloned into pETM-11 vector with N-terminal hexahistidine tag followed by a TEV protease cleavage site. The R235C and Y246D mutations were introduced using the QuickChange method and validated using sequencing. Proteins were overexpressed and purified as previously described (*27*, *31*), using immobilized metal affinity chromatography and size-exclusion chromatography. Briefly, *E. coli* T7 express competent cells, transformed with the GGPPS construct, were grown in terrific broth medium at 37°C until reaching OD_600nm_ = 0.6 and induced at 16°C by adding 0.25 mM isopropyl β-D-1-thiogalactopyranoside (IPTG). Proteins were expressed at 16°C for 16-20 h and harvested by centrifugation (10,000x*g* for 10 min). Cells were suspended in buffer A, containing 50 mM HEPES, pH 7.5, 500 mM NaCl, 5% glycerol, 0.5 mM tris(2-carboxyethyl)phosphine (TCEP), and supplemented with 30 mM imidazole, 1 μg/ml DNase I, 1 mM phenylmethane sulfonyl fluoride (PMSF) and Protease Inhibitor Cocktail Set III (Calbiochem, San Diego, CA, USA). The cells were disrupted in an EmulsiFlex C3 homogenizer (Avestin Inc., Ottawa, ON, Canada). The soluble protein was then recovered by centrifugation at ∼ 32,000x*g* for 45 min at 4 °C. The supernatant was loaded onto a Ni^2+^-NTA column, followed by thorough washing with buffer A, supplemented with 30 mM imidazole to reduce nonspecific protein binding. Next, overexpressed proteins were eluted with buffer A supplemented with 250 mM imidazole. Imidazole was removed using a HiPrep 26/10 desalting column (Cytiva, MA, USA) equilibrated with buffer A, followed by the addition of 6xHis-tagged TEV protease (1 mg TEV protease per 50 mg protein) to the eluant, to remove their 6xHis-tagged, at 4°C overnight. The cleaved proteins were then loaded again onto a Ni^2+^-NTA column with pre-equilibrated with buffer A containing 30 mM imidazole to remove the cleaved 6xHis-tag and TEV protease. The flow-through was collected, concentrated to 3-4 mL, and loaded onto a HiLoad 16/60 Superdex 200 column (Cytiva, MA, USA) equilibrated with 10 mM HEPES, pH 7.5, 100 mM NaCl, 1mM MgCl_2_, 0.5 mM TCEP for final purification. Purified proteins were concentrated and flash-frozen in liquid nitrogen and stored at −80°C.

### SEC-MALS

Experiments were carried out using an analytical SEC column (Superdex 200 10/300 GL), pre-equilibrated with size exclusion buffer containing100 mM NaCl, 10 mM Na-HEPES, 1 mM MgCl_2_, 0.5 mM TCEP, pH 7.5. Samples (2 mg/ml; 50 µl) were injected onto an HPLC, connected to an 8-angle light-scattering detector, followed by a differential refractive index (RI) detector (Wyatt Technology, CA, USA). RI and MALS readings were analyzed with the ASTRA software package (Wyatt Technology, CA, USA) to determine molecular mass.

### Crystallization and structure determination

Initial crystallization screens were performed using purified GGPPS^R235C^ (38 mg/mL) or GGPPS^Y246D^ (38 mg/mL) at 19°C using the sitting drop vapor diffusion method. Initial crystals were obtained in 0.1M sodium cacodylate pH 6.6, 40% MPD, 5% PEG 8000, 0.1M spermine (GGPPS^R235C^) or 1.6M Sodium citrate tribasic dihydrate pH 6.5, 0.1M Trimethylamine hydrochloride (GGPPS^Y246D^). Crystals were cryoprotected with 20% glycerol and immersed in liquid N_2_. Data were collected at 100°K at the European Synchrotron Radiation Facility (ESRF; Grenoble, France). Integration, scaling and merging of the diffraction data were done with the XDS program (*48*). As the data for GGPPS^R235C^ were slightly anisotropic, an ellipsoidal truncation and anisotropic scaling were performed (*49*). The structure was solved by molecular replacement using the programs PHASER (*50*) and PHENIX (*51*) (Table 1). The structure of a single dimer from GGPPS-WT (PDB 2Q80) was used as a search model. Iterative model building and refinement were carried out in PHENIX with manual adjustments using COOT (*52*). Structural illustrations were prepared with PyMOL (https://pymol.org). Atomic coordinates and structure factors for the structure of GGPPS^R235C^ and GGPPS^Y246D^ have been deposited in the Protein Data Bank with accession numbers 9HJS and 9HJZ, respectively.

**Table 1.** Data collection and refinement statistics.

|  | GGPPS<sup>Y246D</sup> | GGPPS<sup>R235C</sup> |
| --- | --- | --- |
|  | PDB 9HJZ | PDB 9HJS |
| **Data collection** |  |  |
| Spacegroup | P622 | P2 |
| Cell dimensions |  |  |
| a,b,c (Å) | 192.73, 192.73, 97.64 | 102.62, 69.08, 154.99 |
| $\alpha,\beta,\gamma$ (°) | 90, 90, 120 | 90, 97.39, 90 |
| Beamline | ESRF ID30A-3 | ESRF ID23-1 |
| Wavelength (Å) | 0.96770 | 0.88560 |
| Resolution (Å) | 2.54 | 2.51 |
| Multiplicity | 12.4 (12.7) | 3.5 (3.5) |
| Completeness (%) | 98.8 (98.7) | 77.2 (20.8) |
| Mean I/$\sigma$(I) | 14.6 (1.1) | 12.5 (1.5) |
| R<sub>meas</sub> (%) | 14.7 (233.0) | 8.9 (150.5) |
| CC<sub>1/2</sub> (%) | 99.8 (52.2) | 99.9 (54.6) |
| **Refinement statistics** |  |  |
| No. reflections (work/free) | 35165/1759 | 56863/1999 |
| Resolution range | 46.85-2.54 | 45.19-2.51 |
| R<sub>work</sub>/R<sub>free</sub> | 0.2227/0.2536 | 0.2272/0.2714 |
| No. atoms |  |  |
| Macromolecules | 4349 | 13402 |
| Ligands | 13 | 186 |
| Solvent | 34 | 81 |
| average B-factor |  |  |
| Macromolecules | 67.8 | 75.0 |
| Ligands | 80.7 | 70.2 |
| Solvent | 61.6 | 62.0 |
| RMSD |  |  |
| Bond lengths (Å) | 0.005 | 0.006 |
| Bond angles (°) | 0.63 | 0.78 |
| Ramachandran outliers (%) | 0 | 0 |

Note: Statistics are based on one crystal per dataset
\*Values in parentheses are for highest-resolution shell.

### nESI analyses

Proteins samples were diluted to 40 µM concentration (per monomer) with 100 mM HEPES pH 7.5, 100 mM NaCl and 0.5 mM TCEP containing either 4mM EDTA or 1 mM MgCl_2_, 40 µM GGPP to populate apo or fully Mg^2+^/GGPP loaded states, respectively. Samples were incubated for 60 minutes on ice, and then buffer exchanged into 150 mM ammonium acetate pH 7.5 by two consecutive passes over centrifugal gel filtration columns (Micro Bio-Spin P-6, 6-kDa cut off; BioRad). Protein concentration was verified by A_280_, and all samples were adjusted to equal 20 µM concentrations. Samples were loaded into in-house prepared golden-coated borosilicate glass capillaries (1B120F-4, World Precision Instruments) and electrosprayed into a Waters Synapt G2Si mass spectrometer. The mass spectrometer was operated in positive sensitivity ion mode over the mass range of 500 – 20,000 m/z and tuned to low ion activation with maximal signal intensity. The ESI capillary and sampling cone were kept at 1.2 kV and 80 V, respectively, source temperature 80°C. Argon was used as the collision gas at 6 ml/min flow. Data were acquired with variable low and high trap collision voltage of 10V and 80V, respectively). Data were smoothed and externally mass recalibrated on caesium iodide clusters in MassLynx 4.2 (Waters) with mass deconvolution in UniDec 6.0.4 (*53*).

### HDX-MS

The H/D exchange reactions (Table 2) were managed by the PAL DHR autosampler (CTC Analytics AG). GGPPS^WT^, GGPPS^R235C^ and GGPPS^Y246D^ variants in 100mM HEPES pH 7.5, 100 mM NaCl and 0.5 mM TCEP were prepared at 10 µM concentration in their apo (buffer supplemented with 1 mM EDTA) or holo states (buffer with 1 mM MgCl_2_ and 10 µM GGPP). The exchange reactions were started by a 10-fold dilution into the identical buffer prepared using D_2_O (pD 7.5). HDX was followed for 20 s, 1 m, 5 m, 20 m, and 2 h (with 20 s, 5 m, and 2 h time points triplicated). The exchange reaction was quenched by adding chilled 1 M glycine-HCl pH 2.3 at a 1:1 (v/v) ratio. Samples were immediately injected onto the LC system maintained at 0°C coupled to an Agilent Infinity II 1260/1290 UPLC (Agilent Technologies). The LC outlet was directly interfaced with the ESI source of timsTOF Pro equipped with PASEF (Bruker Daltonics). Samples were online digested (200 µL/min, 0.4% formic acid in water) on a custom-made pepsin/nepenthesin-2 column (bed volume 66 µL) (*54*) and the peptides desalted on a trap column (SecurityGuard™ ULTRA Cartridge UHPLC Fully Porous Polar C18, 2.1 mm ID; Phenomenex) using the same solvent and flow rate. After three minutes, the peptides were eluted by a water-acetonitrile gradient (10-45% in 6 minutes) on an analytical column (Luna Omega Polar C18, 1.6 µm, 100 Å, 1.0×100 mm; Phenomenex) where they were further separated. The solvents used were A: 0.1% FA in water and B: 0.1% FA, 2% water in ACN. The flow rate on the analytical column was 50 µL/min. The mass spectrometer operated in the MS mode with a 1Hz data acquisition rate. The data were peak picked in DataAnalysis (v. 5.3, Bruker Daltonics), exported to text-based files and further processed using DeutEx software (*55*). Data visualization was performed using MSTools (http://peterslab.org/MSTools/index.php) (*56*) and open-source PyMOL molecular graphics system version 2.6.0 (Schrödinger, LLC). Fully deuterated controls were prepared to correct for back-exchange. For peptide identification, the same LC-MS system as described above was used, but the mass spectrometer was operated in data-dependent MS/MS mode with PASEF active and trapped ion mobility separation enabled. The LC-MS/MS data were searched using MASCOT (v. 2.7, Matrix Science) against a custom-built database combining a common cRAP.fasta (https://www.thegpm.org/crap/), sequences of GGPPS variants and the utilized proteases. Search parameters were set as follows: precursor tolerance 10 ppm, fragment ion tolerance 0.05 Da, decoy search enabled, FDR ˂1%, IonScore ˃20 and peptide length ˃5.

**Table 2.**
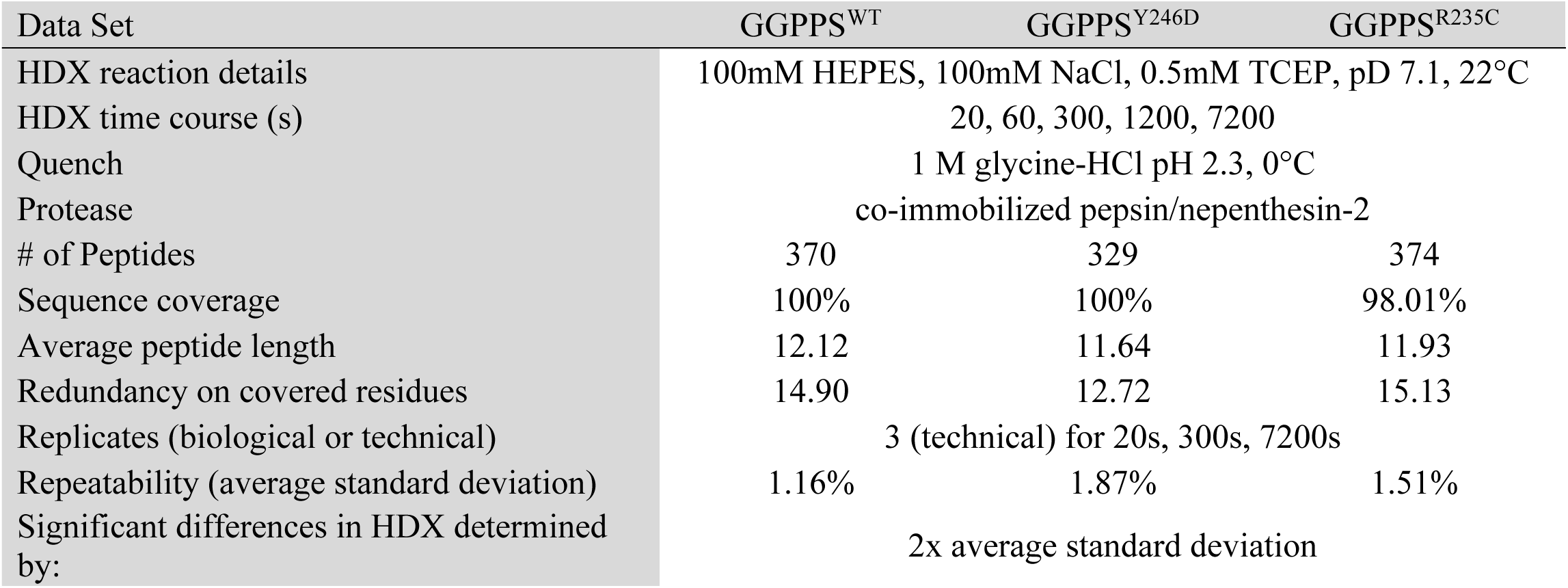
HDX-MS summary statistics.

| Data Set | GGPPS<sup>WT</sup> | GGPPS<sup>Y246D</sup> | GGPPS<sup>R235C</sup> |
| --- | --- | --- | --- |
| HDX reaction details | 100mM HEPES, 100mM NaCl, 0.5mM TCEP, pD 7.1, 22°C |  |  |
| HDX time course (s) | 20, 60, 300, 1200, 7200 |  |  |
| Quench | 1 M glycine-HCl pH 2.3, 0°C |  |  |
| Protease | co-immobilized pepsin/nepenthesin-2 |  |  |
| # of Peptides | 370 | 329 | 374 |
| Sequence coverage | 100% | 100% | 98.01% |
| Average peptide length | 12.12 | 11.64 | 11.93 |
| Redundancy on covered residues | 14.90 | 12.72 | 15.13 |
| Replicates (biological or technical) | 3 (technical) for 20s, 300s, 7200s |  |  |
| Repeatability (average standard deviation) | 1.16% | 1.87% | 1.51% |
| Significant differences in HDX determined by: | 2x average standard deviation |  |  |

### Enzyme kinetics

GGPPS activity was analyzed using a radioactive assay as previously described (*5*). Briefly, 0.02-0.1 μM protein was assayed in a buffer containing 50 mM Tris, 2 mM MgCl_2_, 1 mM TCEP, 5 μg/mL bovine serum albumin, and 0.2% (w/v) Tween 20, pH 7.7. The reaction was initiated by mixing the reaction buffer with [^14^C]IPP (Perkin Elmer, MA, USA) and various FPP concentrations (Sigma Aldrich, MO, USA) at 30 °C. The reactions were quenched by the addition of HCl/methanol (1:4) before reaching 10% of substrate consumption to obtain the initial rates. The product, GGPP (encompassing ^14^C), was extracted by thorough mixing with ligroin, and the product in the ligroin phase was quantitated using a scintillation counter. Data were fit to the Michaelis-Menten equation using GraphPad Prism.

### Fluorescence spectroscopy

All fluorescence experiments were performed in triplicates using an RF-8500 spectrofluorometer (Jasco, Japan) in fluorescence buffer (20 mM Tris-HCl, pH 7.5, 150 mM NaCl, 10 mM β-mercaptoethanol, 0.5 mM MgCl_2_). FRET experiments between GGPPS and MANT-O-GPP were performed using 1 μM of purified GGPPS^WT^ preincubated with 0-40 μM MANT-O-GPP. The competition experiments were performed by preincubation of the enzyme with varying [FPP] (0-100 μM) or [IPP] (0-100 μM) while holding [MANT-O-GPP] constant at 2 μM. Data were plotted using GraphPad Prism.

### Stopped-flow kinetics

Experiments were performed using the RF-8500 spectrofluorometer SFS-852 stopped-flow module (Jasco, Japan). 50 μL of 1 μM the different GGPPS constructs preincubated with 1 μM MANT-O-GPP in fluorescence buffer were mixed with 50 μL of 20 μM FPP in fluorescence buffer at 10 mL/sec, resulting in a dead time of ∼ 4 msec. Fluorescence intensity at 420 nm was measured following excitation at 280 nm. Data were fit to one (GGPPS^WT^) or two-phase (GGPPS^R235C^ and GGPPS^Y246D^) exponential equations with Bio-Kine 32 V4.45.

## Acknowledgments

We acknowledge the European Synchrotron Radiation Facility for the provision of synchrotron radiation facilities, and we would like to thank Dr. Sylvain Engilberge and Dr. Montserrat Soler-López for assistance in using beamlines ID23-1 and ID30A-3, respectively.

## Funding

Israel Science Foundation grant 1653/21 (YH)

Israel Cancer Research Fund RCDA grant 19202 (MG)

Israel Cancer Association grant 20230029 (MG, YH)

Israel Cancer Research Fund Project grant 1289067 (YH, MG)

Bi-National Science Foundation grant 2023190 (YH)

Kahn foundation’s Orion project, Tel Aviv Medical Center, Israel (MG)

Claire and Amedee Maratier Institute for the Study of Blindness and Visual Disorders, Faculty of Medical and Health Sciences, Tel-Aviv University (YH, MG)

MEYS/EU project OP JAK – Photomachines (CZ.02.01.01/00/22_008/0004624) (PM)

CIISB LM2023042 and ERDF “UP CIISB” (CZ.02.1.01/0.0/0.0/18_046/0015974) (PM)

The European Union’s ERA fellowship (101090276) (AK)

## Author contributions

Each author’s contribution(s) to the paper should be listed (we suggest following the CRediT model with each CRediT role given its own line. No punctuation in the initials.

Conceptualization: MG and YH

Methodology: AK, PM, MG, and YH

Investigation: RY, JMP, RMY,AK, PM, MG, and YH

Visualization: AK, PM, MG, and YH

Supervision: MG and YH

Writing – original draft: MG and YH

Writing – review & editing: AK, PM, MG, and YH

## Competing interests

Authors declare that they have no competing interests.

## Data and materials availability

Atomic coordinates and structure factors for the structures of GGPPSR235C and GGPPSY246D have been deposited in the Protein Data Bank under accession codes 9HJS and 9HJZ, respectively. The MS proteomics data have been deposited to the ProteomeXchange Consortium via the PRIDE partner repository with the dataset identifier PXD058943. All other data are available in the main text (Table 2) or the Supplementary Materials (Figs. S1, S2, S3).

## Notes

### Competing Interest Statement

The authors have declared no competing interest.

