## Supplementary information for "A somatic multiple myeloma mutation unravels a mechanism of oligomerization-mediated product inhibition in GGPPS"

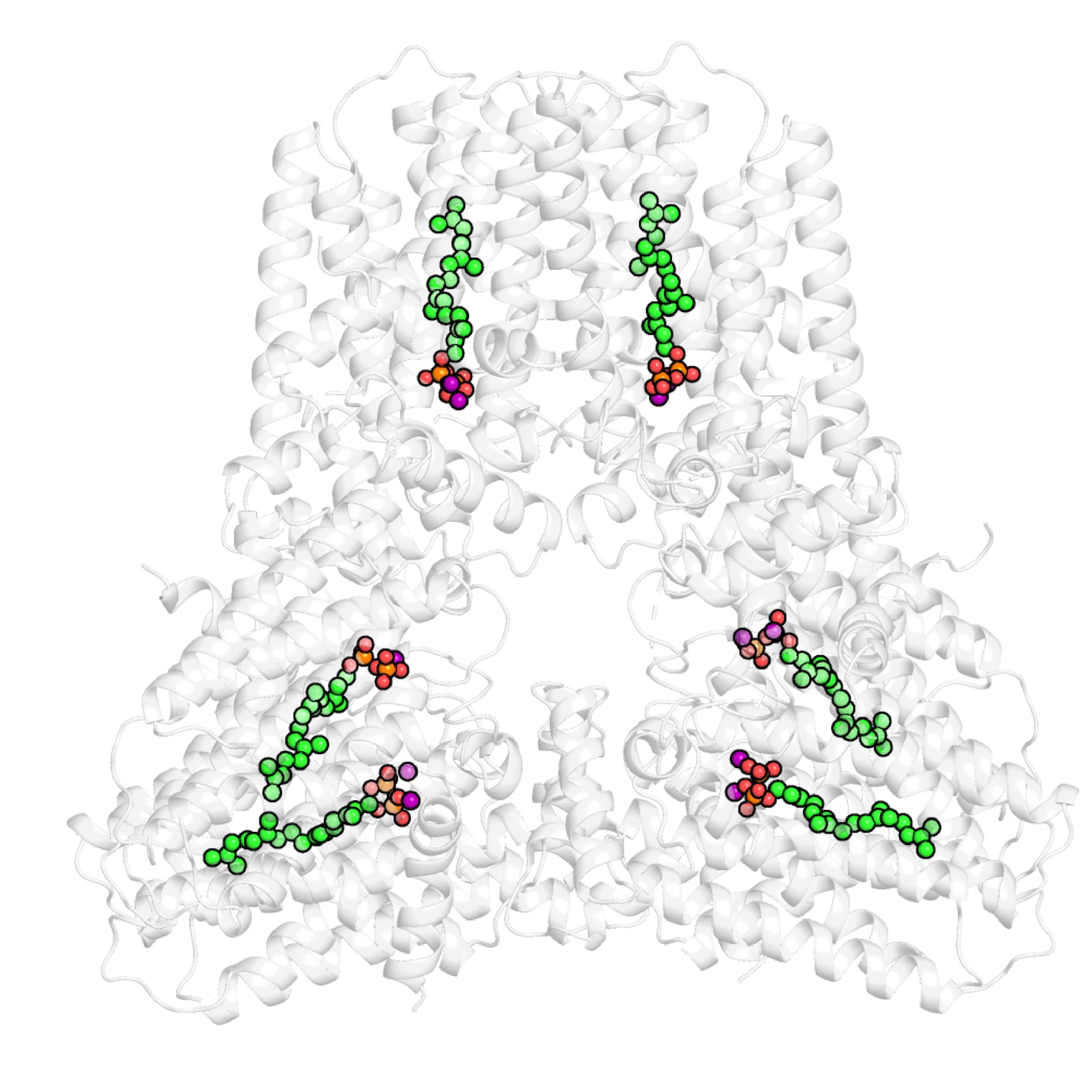

Fig. S1.

**Overall structure of GGPPS^R235C^.** Cartoon representation of the hexameric complex. The GGPP product (green) and magnesium ions (purple) are shown as spheres.

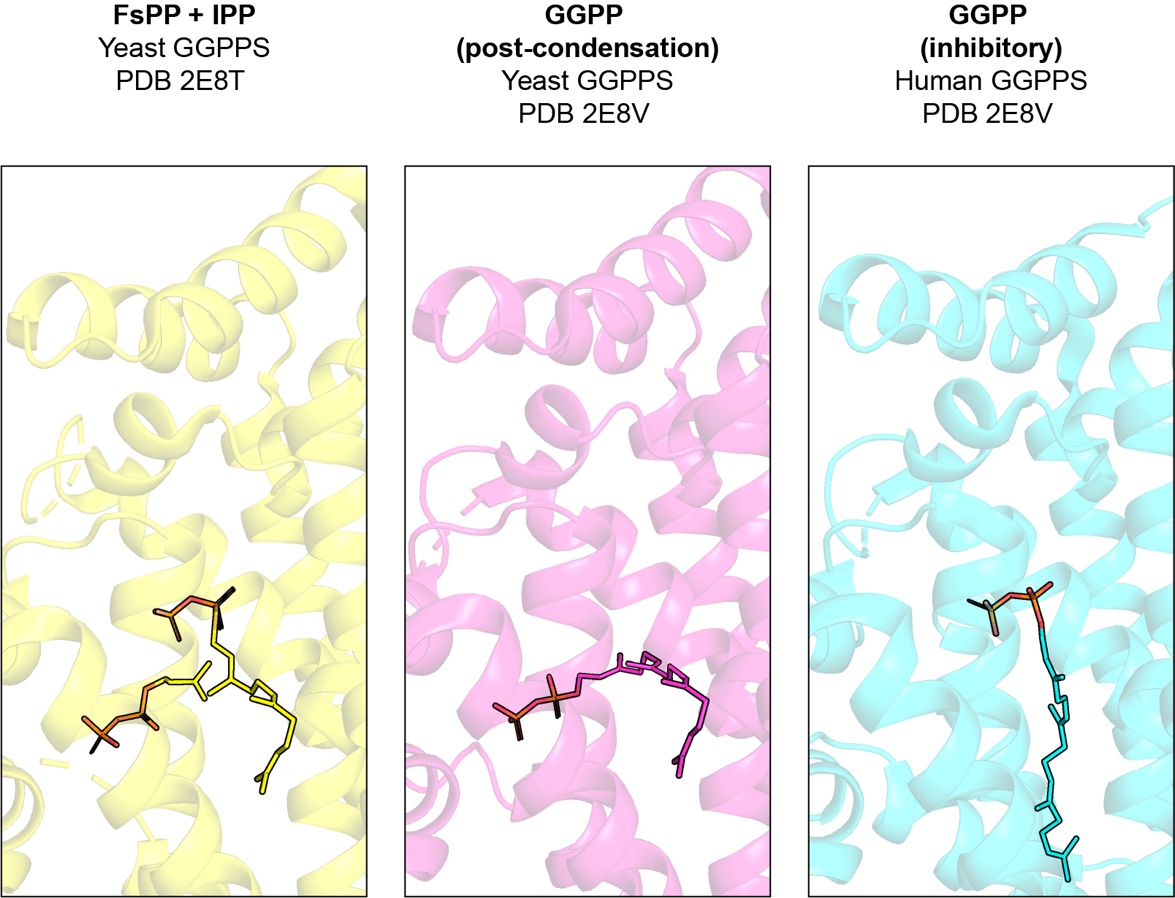

Fig. S2.

**Substrate- and product-bound states of GGPPS.** Blow-out views of differentially occupied active sites of GGPPS. In yeast GGPPS, following the condensation of the substrates (left panel), GGPP occupies a non-inhibitory site (middle panel). In human GGPPS, GGPP translocates to a different conformation, prohibiting the subsequent binding of the FPP substrate (right panel).

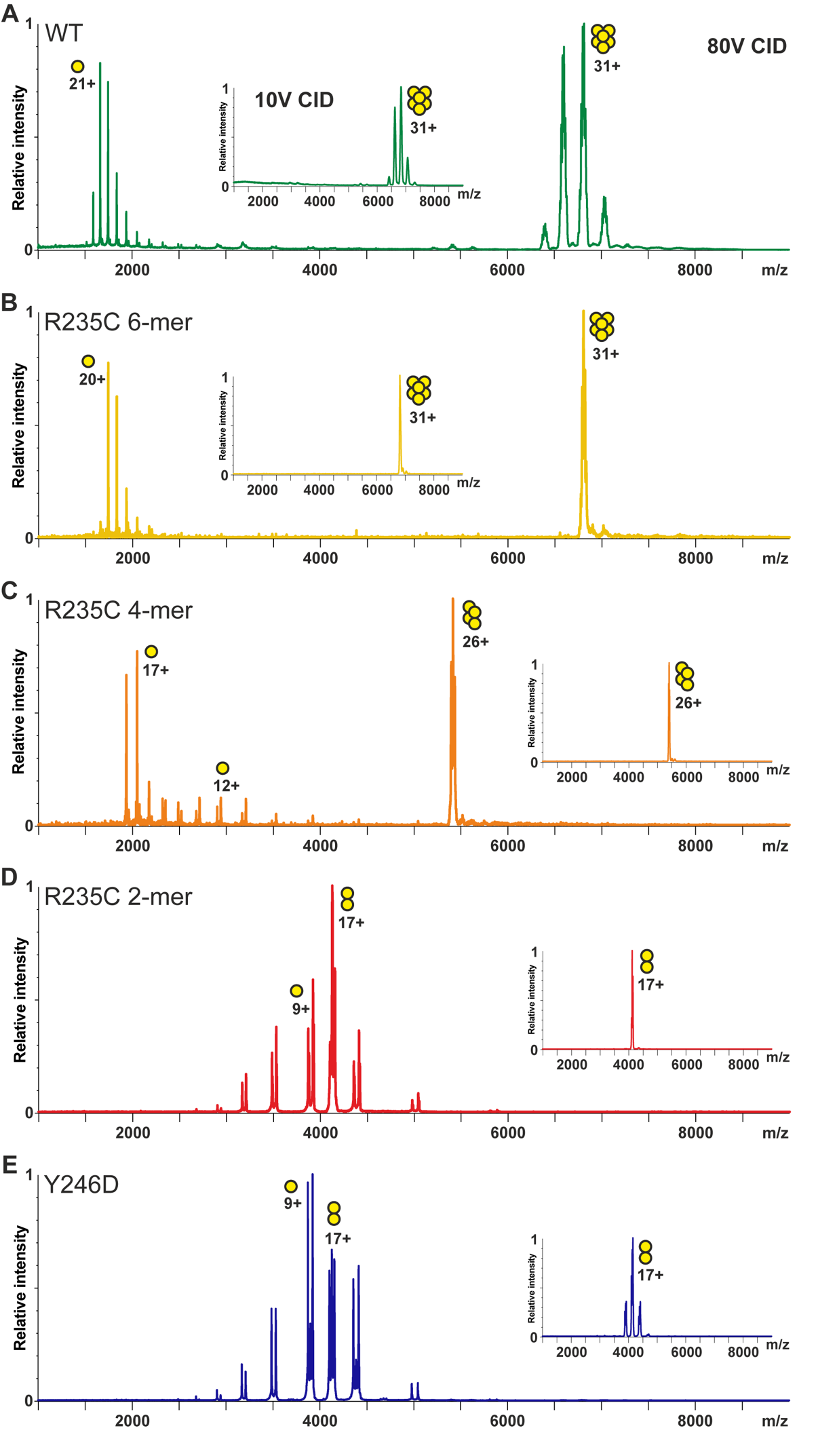

Fig. S3.

**Stability of different GGPPS oligomeric states monitored by nESI. (A-E)** Collision-induced dissociation (CID) at 80V trap voltage shows the ejection of a highly charged unfolded monomeric subunit for GGPPS^WT^ and GGPPS^R235C^ hexamer (A,B). Dimeric species of GGPPS^Y246D^ and GGPPS^R235C^ dissociate into equally charged folded monomers (D,E), while GGPPS^R235C^ tetramer (C) displays a mixture of both modes. Maximum charge states are labeled for all species. Insets show low activation MS spectra of isolated precursor peaks subsequently subjected to 80V CID.
